# Inhibition of TRPV1 and TRPA1 by mosquito and mouse saliva

**DOI:** 10.1101/2020.05.12.090985

**Authors:** Tianbang Li, Sandra Derouiche, Yuya Sakai, Daisuke Uta, Seiji Aoyagi, Makoto Tominaga

## Abstract

Arthropods are the largest group of living organisms and among them, mosquitoes spread parasites and viruses causing deathly diseases. Mosquito’s painless piercing is due to the thinness of their fascicle, but mosquito saliva might also contribute to it. We found that mosquito and mouse saliva inhibit TRPV1 and TRPA1 channels, either heterologously expressed in HEK293T cells or endogenously expressed in native mouse sensory neurons. We have also identified sialorphin as a candidate antinociceptive peptide as it showed similar effects on TRPV1 and TRPA1. Finally, we confirmed the antinociceptive effects of saliva and sialorphin *in vivo* by observing decreased pain-related behaviors in mice coinjected with these substances. Similar inhibitory effects of mosquito and mouse saliva suggest that the antinociceptive effects of saliva are universal, which could explain why many animals including humans often lick their wounds. These findings would lead to the development of novel anti-nociceptive agents.

## Introduction

Arthropods are the largest group of living organisms. They attack other organisms by biting, stinging, piercing and sucking. Among the various arthropods which impact human health by feeding on living hosts, piercing by mosquitoes (*Figure 1A*) spreads parasites and viruses, some of which have been reported to cause the highest number of deaths annually, including malaria, Dengue fever and West Nile fever (Tolle, 2009).

**Figure 1.**
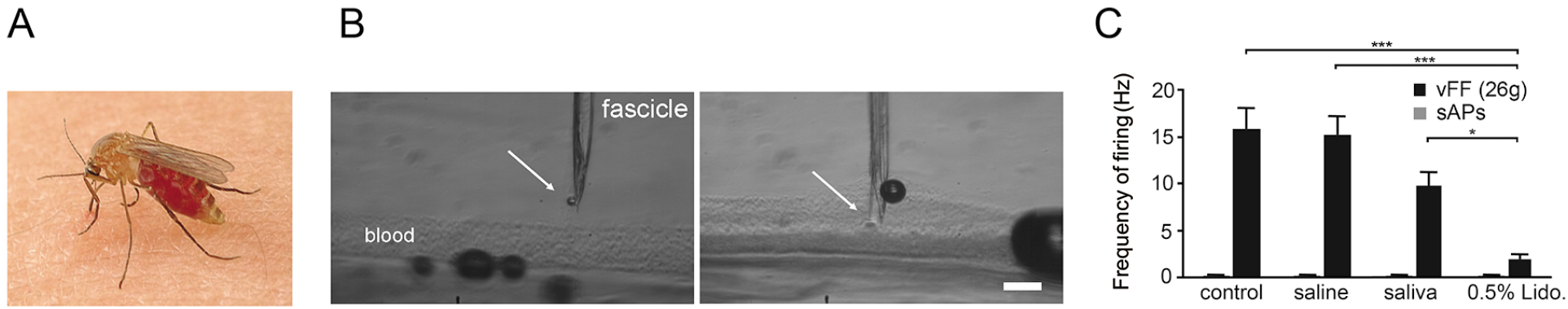
Inhibition of firing of spinal cord neurons by mosquito saliva. (*A*) Photograph of *Culex pipiens pallens* on human skin. (*B*) Mosquito saliva discharge into artificial blood vessels was fabricated inside the skin using Micro-Electro-Mechanical Systems technology. Arrows indicate the saliva discharge at the tip of fascicle. Scale bar: 100 μm. (*C*) Inhibition of von Frey filament (vFF) evoked neuronal firing frequency in rat spinal dorsal horn neurons by 0.5 % lidocaine (Lido) or a 10-fold dilution of mosquito (*Culex pipiens pallens*) saliva. n = 7 each. sAPs: spontaneous action potentials. *** p < 0.001, * p < 0.05 by one-way ANOVA followed by a Bonferroni post hoc test.

The mosquito fascicle, which is composed of six stylets, is approximately 70 μm in diameter, being far thinner than commercial medical needles of which diameter are usually over several hundred μm (Suzuki, 2008). One of the reasons why mosquito piercings are painless is the thinness of their fascicle, and the potential for fascicles to avoid pain receptors (free nerve endings) becomes higher as they become thinner. Over their long process of evolution, mosquitoes have adapted by dividing the needle into a fascicle of six stylets (Heinemann, 2008), and mosquitoes can move these stylets with great dexterity, which also contributes to the painless piercing (Clement, 2000). Saliva is discharged through cooperative movement of the stylets even before they reach the blood supply (*Figure 1B*). Once they reach the blood source, the cooperative motion of the stylets ceases and saliva is frequently discharged as they suck the blood (Izumi, Suzuki, Aoyagi, & Kanzaki, 2011).

Although the microneedle properties of the fascicle are currently believed to explain the painless piercings by mosquitoes, it is possible that mosquito saliva might also contribute to this lack of pain. Mosquito saliva is a complex of proteins, many of which have unknown functions, but some allow the mosquito to acquire a blood meal from its host by circumventing vasoconstriction, platelet aggregation, coagulation, and inflammation or hemostasis (Sun, McNicol, James, & Peng, 2006). Mosquito saliva also contains proteins that are immunogenic to humans and cause allergic responses, as we often experience. Given these properties, we hypothesized that mosquito saliva also contains antinociceptive substances that contribute to their painless piercings.

## Results

### Inhibition of firing of spinal cord neurons by mosquito saliva

To verify the possibility of antinociceptive effects of mosquito saliva, we first examined the activity of mosquito saliva in rat spinal cord neurons upon mechanical stimulation (Uta et al., 2019). In the second layer of spinal dorsal horn neurons of rats, von Frey filament (26 g)-evoked firing frequencies looked reduced upon injection of a 10-fold dilution of mosquito saliva although statistical significance was not obtained (15.2 ± 2.2 Hz and 9.8 ± 1.7 Hz for saline and saliva, respectively; p = 0.30; *Figure 1C*), suggesting the possible antinociceptive effects of mosquito saliva. We confirmed drastic reduction of the firing frequencies upon injection of 0.5 % lidocaine (1.9 ± 0.6 Hz; p < 0.001 vs. control, p < 0.001), which inhibits activation of voltage-gated Na+ channels (Hille, 1966). Spontaneous firings were hardly observed.

### Inhibition of TRPV1- and TRPA1-mediated currents by mosquito or mouse saliva

Under normal conditions, mosquito saliva should act on the sensory neurons innervating the epidermis, and there are several ion channels expressed in the sensory nerve endings which can detect noxious stimuli, among these are the TRP channels. Given this, we next examined the effects of mosquito saliva on TRP channel function, focusing on human TRPV1 and human TRPA1 (Julius, 2013), both of which are reported to be activated by mechanical stimuli which is supposed to function during mosquito piercing in addition to noxious chemicals or high temperature in HEK293T cells (Fujita et al., 2018; Moore & Liedtke, 2017). We held the membrane potential at 0 mV and applied voltage ramp-pulses from −100 to +100 mV every 5 sec for the recording of TRPV1 currents. Interestingly, diluted mosquito saliva significantly inhibited the human TRPV1 currents activated by 50 nM capsaicin in a dose-dependent and reversible manner without apparent voltage dependency observed in the current-voltage (I-V) curves (−2.4, 7.1, 21.1 and 31.8 % inhibition by 300-, 100-, 20- and 10-fold dilution of saliva, respectively, at +100 mV; p < 0.05 vs. control at 10-fold dilution; *Figure 2A and B*). Furthermore, we examined the effects of mosquito saliva on human TRPA1 channel in which we used citronellal because the currents activated by allyl isothiocyanate (AITC) or cinnamaldehyde were hardly stabilized. And we held the membrane potential at −60 mV and applied voltage ramp-pulses similar to the TRPV1 current measurement. Mosquito saliva inhibited human TRPA1 currents activated by 500 μM citronellal in a dose-dependent and reversible manner without apparent voltage dependency observed in the I-V curves while TRPA1-mdediated currents were reduced gradually even in the control condition (52.8, 56.5, 63.4 and 55.4 % inhibition by 300-, 100-, 20- and 10-fold dilution of saliva, respectively, at −60 mV; p < 0.01 vs. control at 10-fold dilution; *Figure 2C and D*). In order to evaluate whether these effects of saliva are common to other animals, we also examined the effects of mouse saliva on mouse TRPV1 and TRPA1 channels. Similar to the mosquito saliva, even a 100-fold dilution of mouse saliva significantly inhibited both mouse TRPV1- and TRPA1-mediated current responses activated by capsaicin (20 nM) or citronellal (500 μM), respectively (97.4 ± 0.9 and 90.5 ± 2.2 % of the values before application in control and saliva, respectively, p < 0.01 for TRPV1, and 86.5 ± 2.6 and 53.0 ± 2.8 % of the values before application in control and saliva, respectively, p < 0.001 for TRPA1; *Figure 2E-H*).

**Figure 2.**
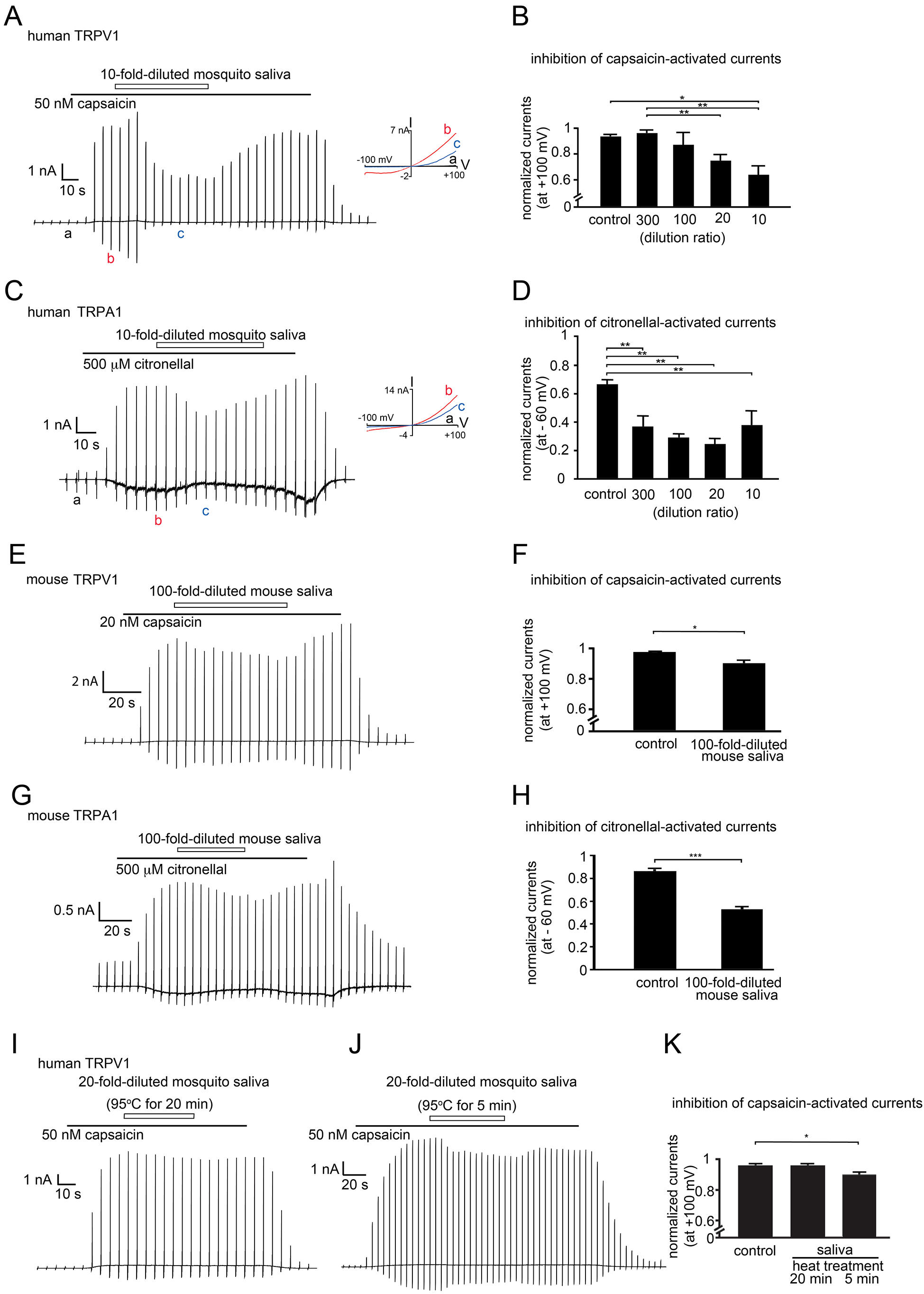
Inhibition of TRPV1- and TRPA1-mediated currents by mosquito or mouse saliva. (*A*) Representative trace of inhibition of capsaicin (50 nM)-activated human TRPV1 currents by a 10-fold dilution of mosquito saliva. Holding potential was 0 mV with ramppulses (−100 ~ +100 mV, 300 ms) applied every 5 sec. (inset) current-voltage (I-V) curve at the points indicated by a, b and c in the trace. (*B*) Comparison of the normalized currents at +100 mV with saline (control) or mosquito saliva at different dilution ratios. n = 8, 6, 5, 9 and 7 for control, and 300-, 100-, 20- and 10-fold dilution, respectively. * p < 0.05, ** p < 0.01 by one-way ANOVA followed by a Bonferroni post hoc test. (*C*) Representative trace of inhibition of citronellal (500 μM)-activated human TRPA1 currents by a 10-fold dilution of mosquito saliva. Holding potential was −60 mV with ramp-pulses (−100 ~ +100 mV, 300 ms) applied every 5 sec. (inset) I-V curve at the points indicated by a, b and c in the trace. (*D*) Comparison of the normalized currents at −60 mV with saline (control) or mosquito saliva at different dilution ratios. n = 6, 6, 6, 6 and 5 for control, and 300-, 100-, 20- and 10-fold dilution, respectively. ** p < 0.01 by one-way ANOVA followed by a Bonferroni post hoc test. (*E*) Representative trace of inhibition of capsaicin (20 nM)-activated mouse TRPV1 currents by a 100-fold dilution of mouse saliva. Holding potential was 0 mV with ramppulses (−100 ~ +100 mV, 300 ms) applied every 5 sec. (*F*) Comparison of the normalized currents at +100 mV with saline (control, n = 11) or mouse saliva (n = 7). * p < 0.05 by Student’s *t* test. (*G*) Representative trace of inhibition of citronellal (500 μM)-activated mouse TRPA1 currents by a 100-fold-dilution of mouse saliva. Holding potential was −60 mV with ramppulses (−100 ~ +100 mV, 300 ms) applied every 5 sec. (*H*) Comparison of the normalized currents at −60 mV with saline (control, n =5) or mouse saliva, n = 6). * p < 0.05 by Student’s *t* test. (*I and J)* Representative traces of inhibition of the capsaicin (50 nM)-activated human TRPV1 currents by a 20-fold dilution of mosquito saliva treated with 20 *(I)* or 5 *(J)* min of 95oC heat. Holding potential was 0 mV with ramp-pulses (−100 ~ +100 mV, 300 ms) applied every 5 sec. *(K)* Comparison of the normalized currents at +100 mV with heat-treated mosquito saliva for 20 or 5 min. n = 8, 5 and 5 for control, 20-fold diluted mosquito saliva with 20 min or 5 min heat treatment, respectively. * p < 0.05 by one-way ANOVA followed by a Bonferroni post hoc test.

### Loss of inhibition of TRPV1 currents by heated mosquito saliva

We then sought to determine which classes of substances found in saliva have the ability to inhibit TRPV1 and TRPA1. Heat treatment of mosquito saliva at 95°C for 20 min caused a loss of the TRPV1-inhibiting effects while treatment at 95°C for only 5 min did not (94.0 ± 1.4%, 95.0 ± 1.1% and 88.2 ± 1.6% of the values before application for control, 20 and 5 min treatments, respectively, p < 0.05 for 5 min vs. control; *Figure 2I-K*), suggesting the involvement of peptides in the antinociceptive effects of saliva.

### Inhibition of human TRPV1- and TRPA1-mediated currents by sialorphin

It has been reported that an endopeptidase, sialorphin (*Figure 3A*), which is widely found in saliva, has some antinociceptive effects when applied *in vivo* (Rougeot et al., 2003). Accordingly, we examined the effects of sialorphin on TRP channel functions. Interestingly, 10 μM of sialorphin significantly inhibited human TRPV1- and TRPA1-mediated currents in a dose-dependent and reversible manner without apparent voltage dependency observed in the I-V curves (for human TRPV1: 94.0 ± 1.4, 97.6 ± 1.0, 86.4 ± 3.1, 80.4 ± 3.3 and 72.1 ± 8.8 % of the values before application for control, 0.3, 1, 3 and 10 μM sialorphin, respectively, p < 0.01 vs. control at 10 μM, and for human TRPA1: 66.6 ± 3.3, 38.7 ± 11.8, 47.5 ± 10.1, 37.4 ± 5.0 and 7.6 ± 1.1 % of the values before application for control, 0.3, 1, 3 and 10 μM sialorphin, respectively, *Figure 3B-E*), a phenomenon identical in both mosquito and mouse saliva. This implicated sialorphin as a strong candidate for causing the antinociceptive effects of saliva.

**Figure 3.**
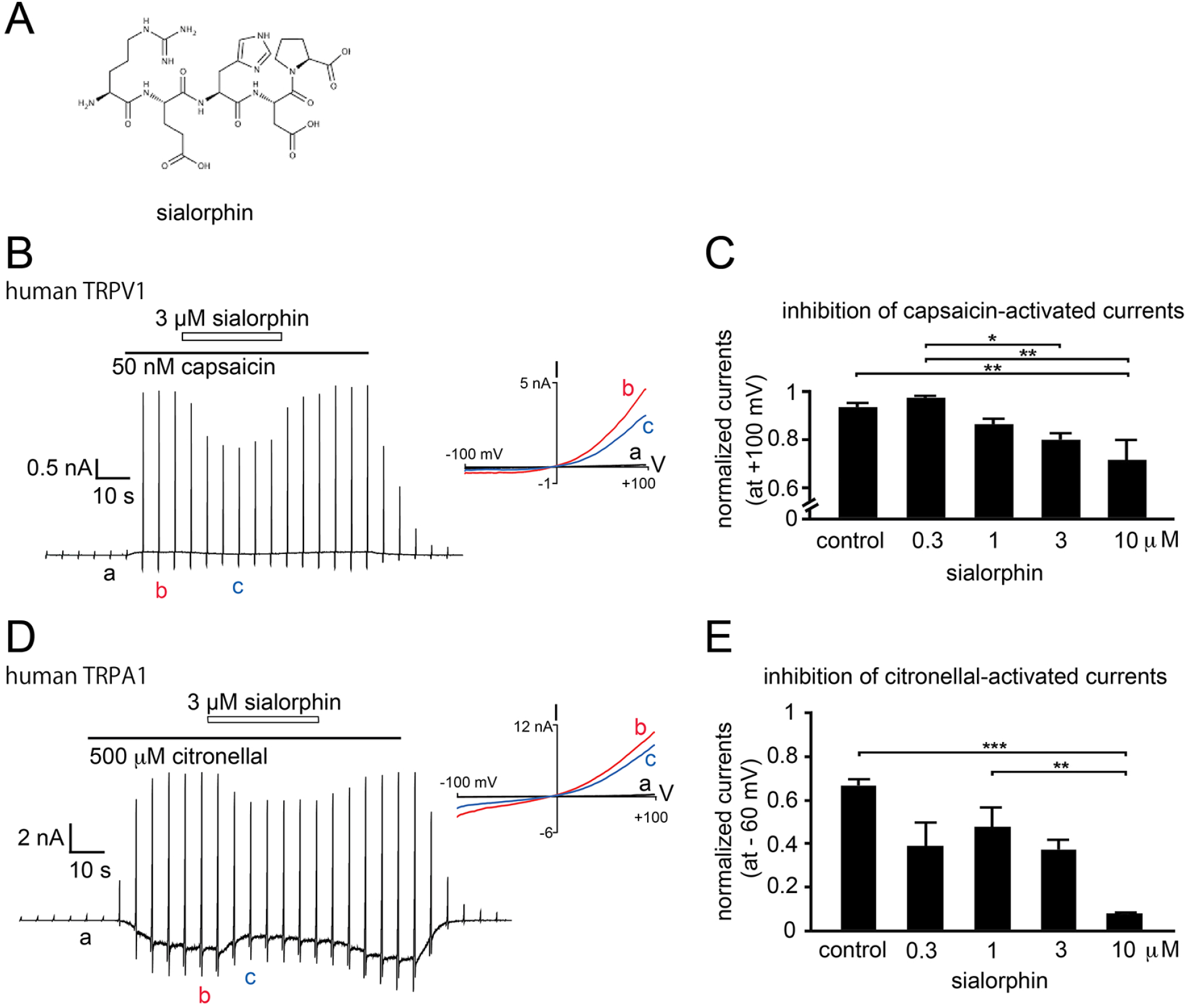
Inhibition of human TRPV1- and TRPA1-mediated currents by sialorphin. (*A*) Structure of sialorphin. (*B*) Representative trace of inhibition of capsaicin (50 nM)-activated human TRPV1 currents by sialorphin (3 μM). Holding potential was 0 mV with ramp-pulses (−100 ~ +100 mV, 300 ms) applied every 5 sec. (inset) I-V curve at the points indicated by a, b and c in the trace. (*C*) Comparison of the normalized currents at +100 mV with saline (control) or sialorphin at different concentrations. n = 8, 6, 6, 8 and 5 for control, and 0.3, 1, 3 and 10 μM sialorphin, respectively. * p < 0.05, ** p < 0.01 by one-way ANOVA followed by a Bonferroni post hoc test. (*D*) Representative trace of inhibition of citronellal (500 μM)-activated human TRPA1 currents by sialorphin (3 μM). Holding potential was −60 mV with ramp-pulses (−100 ~ +100 mV, 300 ms) applied every 5 sec. (inset) I-V curve at the points indicated by a, b and c in the trace. (*E*) Comparison of the normalized currents at −60 mV with saline (control) or sialorphin at different concentrations. n = 5, 5, 5, 5 and 5 for control, and 0.3, 1, 3 and 10 μM sialorphin, respectively. * p < 0.05, ** p < 0.01 by one-way ANOVA followed by a Bonferroni post hoc test.

### Mosquito saliva-induced inhibition of TRPV1- or TRPA1-mediated responses in mouse DRG neurons

We next aimed to determine if similar inhibition is observed in mice sensory neurons. Mosquito saliva diluted 10-fold inhibited the capsaicin-activated currents in native sensory neurons from mouse dorsal root ganglia (DRG) which consist of different peripheral sensory neurons including ones that express TRPV1 or TRPA1 (Takayama, Uta, Furue, & Tominaga, 2015) (*Figure 4A*). The capsaicin-induced increase in intracellular Ca_2+_ concentrations ([Ca_2+_]_i_) also looked slightly reduced by a 20-fold dilution of mosquito saliva, although this difference was not statistically significant (*Figure 4B*). However, the population of capsaicin-sensitive neurons was significantly smaller in the presence of a 20-fold dilution of mosquito saliva (34.0 ± 1.5, 23.1 ± 3.3 and 45.1 ± 6 % for + saline, + saliva and + sailorphin, respectively, p < 0.05 for capsaicin + saliva vs. capsaicin + saline) (*Figure 5C*). Interestingly, sialorphin (3 μM) actually enhanced the capsaicin-induced [Ca_2+_]_i_ increase, suggesting some interaction between the two compounds. In contrast, neither mosquito saliva nor sialorphin affected the AITC (300 μM)-induced [Ca_2+_]_i_ increases (*Figure 5D*), although the population of AITC-sensitive neurons seemed smaller (*Figure 4E*), suggesting possible inhibitory effects of saliva and sialorphin on TRPA1-mediated responses. These results suggested that mosquito saliva has the potential to inhibit TRPV1 and TRPA1 in sensory neurons and that sialorphin could be one of the components contributing to this inhibition.

**Figure 4.**
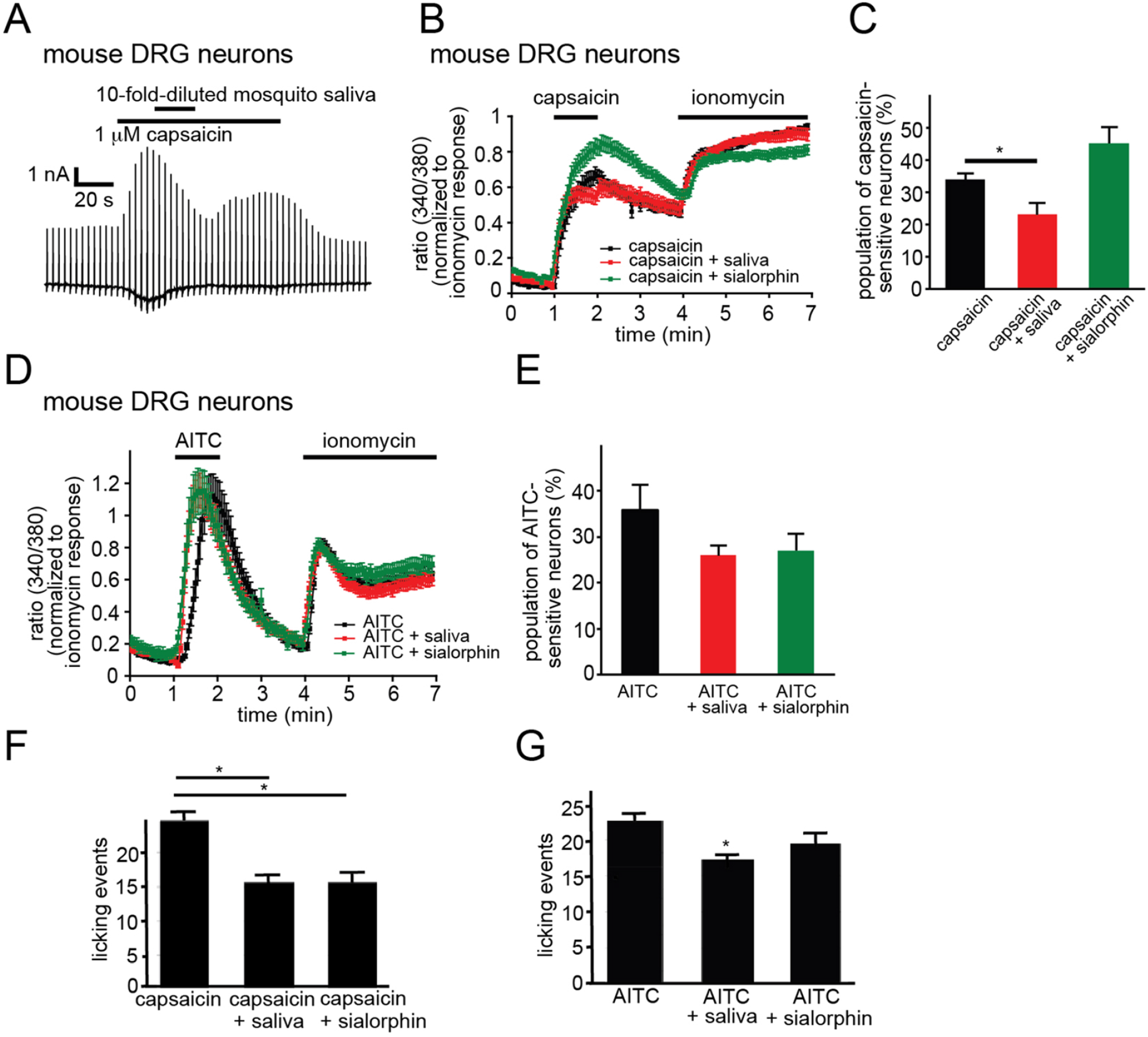
Mosquito saliva-induced inhibition of TRPV1- or TRPA1-mediated responses in mouse DRG neurons and pain-related behaviors in mice. (*A*) Representative trace of inhibition of capsaicin (1 μM)-activated currents by a 10-fold dilution of mosquito saliva in a mouse DRG neuron. Similar results were observed in 3 additional cells. Holding potential was −60 mV with ramp-pulses (−100 ~ +100 mV, 300 ms) applied every 3 sec. (*B*) Mean changes in intracellular Ca_2+_ concentrations (Fura-2 340/380 ratios) in mouse DRG neurons treated with capsaicin (50 nM) + saline (n = 32, black), AITC + 20-fold-diluted mosquito saliva (n = 26, red) or sialorphin (3 μM, n = 67, green), respectively. (*C*) Comparison of the population of capsaicin-sensitive neurons. * p < 0.05 by one-way ANOVA followed by a Bonferroni post hoc test. (*D*) Mean changes in Fura-2 ratios in mouse DRG neurons for AITC (300 μM) + saline (n = 56, black), AITC + 20-fold-diluted mosquito saliva (n = 58, red) or AITC + sialorphin (3 μM, n = 38, green), respectively. (*E*) Comparison of the population of AITC-sensitive neurons. (*F*) Comparison of the total licking events in mice injected with capsaicin + saline (control, 24.8 times, n = 10), + 2-fold-diluted saliva (middle, 15.7 times, n = 11) or + sialorphin (right, 15.6 times, n = 11). * p < 0.05 by one-way ANOVA followed by a Bonferroni post hoc test. (*G*) Comparison of total licking events in mice injected with AITC + saline (control, 22.6 times, n = 14), + 2-fold-diluted saliva (middle, 17.3 times, n = 15) or + sialorphin (right, 19.5 times, n = 15). * p < 0.05 by Student’s t-test.

### Mosquito saliva-induced inhibition of capsaicin- or AITC-induced pain-related behaviors in mice

In order to confirm the antinociceptive effects of saliva and sialorphin *in vivo*, we evaluated pain-related licking behaviors in mice for 5 min after injection of 10 μl of 1 μM capsaicin or 10 μl of 100 mM AITC into their hind paw. Co-injection with either mosquito saliva (10 μl of 2-fold-diluted saliva) or sialorphin (10 μl of a 10 μM solution) into the hind paws of mice caused a significant reduction in licking behaviors caused by capsaicin compared to saline (*Figure 4F*). There were less effects of mosquito saliva (10 μl of 2-fold-diluted saliva) or sialorphin (10 μl of a 10 μM solution) on AITC-induced licking behaviors (*Figure 4G*).

## Discussion

From the data presented above, we conclude that mosquito and mouse saliva exhibit antinociceptive effects through inhibition of TRPV1 and TRPA1, and that sialorphin is a candidate substance contained in saliva that may cause these effects. Mosquitoes seem to use this saliva-induced inhibition of TRPV1 and TRPA1 and skillful movement of their thin fascicle in concert to effectively enable painless piercing. Given the results that both mosquito and mouse saliva showed similar inhibitory effects on TRPV1 and TRPA1, this suggests that the antinociceptive effects of saliva are universal, which could explain why many animals including humans often lick their wounds. The fact that sialorphin is known to be contained in most of saliva supports this concept.

Although the inhibition of TRPV1 or TRPA1 by mosquito saliva is obvious in HEK293T cells (*Figure 2*), we did not see its clear effects in firings of the spinal cord neurons in rats (*Figure 1*), currents or [Ca_2+_]_i_ changes in mouse mRG neurons (*Figure 4A-E*) or pain-related behaviors in mice (*Figure 4F and G*). It could be partly because a lot of TRPV1 or TRPA1 proteins are expressed in HEK293T cells, which makes it easier to see the effects. In the case of behaviors, some other substances or pathways might be necessary to observe the effects on licking behaviors. However, the possibility of involvement of other substances in saliva effects seems unlikely since the effects were not so different between mosquito saliva and sialorphin (*Figure 4*).

We observed further increases of capsaicin-induced [Ca_2+_]_i_ in mouse DRG neurons in the presence of sialorphin (*Figure 4B*). Although we have currently no idea explaining this result, there might be some interaction of sialorphin with capsaicin, but not with AITC, which could be investigated further in the future. However, the possible interaction won’t interfere with the antinociceptive effects of sialorphin since sialorphin did reduce the licking behaviors (*Figure 4F*). In any case, the findings in our study could lead to the development of novel analgesic agents.

## Material and methods

### Animals

Adult mosquito of *Culex pipiens pallens* were from Research & Development Laboratory, Dainihon Jochugiku Co., Ltd. (Osaka, Japan). Mouse experiments were conducted with 6-to 8-week-old C57BL/6NCr male mice. Rat experiments were conducted with 6-to 8-week old of Sprague–Dawley rats. All the animal experiments were performed in accordance with the guidelines of National Institute for Physiological Sciences, University of Toyama and the National Institutes of Health (NIH).

### Chemicals

Capsaicin, citronellal, allyl isothiocyanate (AITC) and carbachol were purchased from Sigma-Aldrich (United States).

### Taking saliva from mosquitoes

After anesthetized with diethyl ether, the mosquito heads with salivary grands were excised using 50 female mosquitoes, followed by homogenized in 800 μL of 0.9% saline, and centrifuged at 15,000 rpm for 15 minutes. The supernatant solution of 700 μL obtained here was used as the mosquito saliva solution. If the larger amount was necessary, the abovementioned process was repeated for each 50 mosquitoes.

### Taking saliva from mice

Mice were initially anesthetized by inhaling isoflurane, followed by a subsequent intraperitoneal injection of 450 μL of pentobarbital (5 mg/ml in PBS) to ensure a complete anesthesia. Mouse neck skin was removed and both salivary glands were exposed for the injection of carbachol. 20 μL of carbachol (50 μM) was injected into each submandibular gland twice discontinuously for stimulating the saliva secretion. Saliva was directly collected with a pipette into an Eppendorf tube and kept at −20 °C until use.

### Cell Culture

The human embryonic kidney 293T (HEK293T) cells were cultured in Dulbecco’s modified Eagle Medium (Wako) supplemented with 10% fetal bovine serum (BioWest), penicillin-streptomycin (50 mg/mL and 50 units/mL, respectively, Gibco) and GlutaMAX (2 mM, Gibco). For transient transfection of HEK293T cells, 1 μg of plasmid DNA in pcDNA3.1 (+) and 0.1 μg pGreen-Lantern 1 vector were transfected into HEK293T cells using Lipofectamine reagent and Plus reagent (Invitrogen). In the case of transfection for calcium-imaging, 0.1 μg pCMV-DsRed vector was transfected instead of pGreen-Lantern 1 vector. All these components were dissolved in OPTI-MEM medium (1X, Gibco). After incubation for 3-4 h, HEK293T cells were reseeded on 12-mm cover slips (Matsunami) and further incubated at 33°C in 5% CO_2_.

### Isolation of mouse DRG neurons

Following anesthesia with isoflurane, the DRG was separated from the L4 to L6 of mice after perfusion with 10 mL ice-cold artificial cerebrospinal fluid (aCSF: 124 mM NaCl, five mM KCl, 1.2 mM KH_2_PO_4_, 1.3 mM MgSO_4_, 2.4 mM CaCl_2_, ten mM glucose, 24 mM NaHCO_3_, equilibrated with 95% O_2_ and 5% CO_2_ for 1 h on ice). The tissues were incubated with 725 μg collagenase type IX (lot# SLBG3258, Sigma-Aldrich) in 250 μL Earle’s balanced salt solution (Sigma-Aldrich) containing 10% FBS (as above), MEM vitamin solution (1:100, Sigma-Aldrich), penicillin/streptomycin (1:200, Life Technologies) and GlutaMax (1:100, Life Technologies) at 37°C for 25 min. Next, the DRG neurons were mechanically separated by 10 to 20 cycles of pipetting using a small diameter Pasteur pipette and filtered through a 40 μm cell strainer (BD Falcon, United States). The isolated neurons were placed on 12-mm diameter coverslips (Matsunami, Japan) with 40 μL aCSF and used for experiments within four h of isolation, maintaining them at room temperature in a 95% O_2_ and 5% CO_2_ humidified chamber.

### Calcium Imaging and Electrophysiology

Both calcium-imaging and whole-cell patch-clamp recording experiments were performed 18-30 h after transfection. The extracellular standard bath solution contained 140 mM NaCl, 5 mM KCl, 2 mM MgCl_2_, 2 mM CaCl_2_, 10 mM HEPES and 10 mM glucose at pH 7.4, adjusted with NaOH. Cytosolic-free Ca_2+_ concentrations were measured with Fura-2 (Molecular Probes, Invitrogen Corp). Fura-2-AM (5 μM) was loaded 1 h before recording, and it was excited at 340/380 nm with emission at 510 nm. Fura-2 fluorescence was recorded with a CCD camera, Cool Snap ES (Roper Scientific/Photometrics). Data were acquired using imaging processing software (IPlab Scanalytic) and analyzed with ImageJ (NIH). For whole-cell patch-clamp recording, the intracellular pipette solution contained 140 mM KCl, 5 mM EGTA, and 10 mM HEPES at pH 7.4 adjusted with KOH. Recording started 2 to 3 min after making a whole-cell configuration to achieve steady state. The data from whole-cell patch-clamp recordings were acquired at 10 kHz and filtered at 5 kHz for analysis (Axopatch 200B amplifier with pCLAMP software, Molecular Devices). Membrane potential was clamped at 0 or −60 mV and voltage ramp-pulses from −100 to +100 mV (300 ms) were applied every 5 s.

### *In vivo* extracellular recording in rat

The methods used for the *in vivo* patch-clamp recording of SG neurons were similar to those described previously (2). In brief, the rats were anesthetized with urethane (1.5 g/kg, i.p.), which produces a long-lasting steady level of anesthesia and does not require the administration of additional doses except in few cases. A thoracolumbar laminectomy was performed to expose the L1–L6 vertebrae, followed by placing the animal in a stereotaxic apparatus. Next, the dura was removed and the arachnoid membrane was cut to create a large window for a tungsten microelectrode. The surface of the spinal cord was irrigated with Krebs solution equilibrated with 95% O_2_ and 5% CO_2_ (10–15 mL/min) and containing 117 mM NaCl, 3.6 mM KCl, 2.5 mM CaCl_2_, 1.2 mM MgCl_2_, 1.2 mM NaH_2_PO_4_, 11 mM glucose, and 25 mM NaHCO_3_ (pH = 7.4) at 37±1°C. Extracellular single-unit recordings of superficial dorsal horn (lamina I and II) neurons were performed as described previously (3). Recordings were obtained from the superficial dorsal horn neurons at a depth of 20-150 μm from the surface. These cells were within the superficial dorsal horn and assessed from slices obtained from the same spinal level of same-age mice. Unit signals were acquired with an amplifier (EX1; Dagan Corporation, Minneapolis, MN, U.S.A.). The data were digitized with an analog-to-digital converter (Digidata 1400A; Molecular Devices, Union City, CA, U.S.A.) stored on a personal computer with a data acquisition program (Clampex software, version 10.2, Molecular Devices) and analyzed with Clampfit software (version 10.2, Molecular Devices). We searched the area on the skin where touch (with a cotton wisp) or noxious pinch (with forceps) stimulus produced a neural response. Mechanical stimulus was applied by skin folding using a fine von Frey filament at a bending force of 255 mN. The stimuli were applied for 10 s to the ipsilateral hind limb at the maximal response point of the respective receptive area.

### Pain-related behavior test

Mice were twice-handled gently for 20 min every 48 h before the behavior test. Mice were injected with capsaicin or AITC with or without saliva or sialorphin (total 10 μL) into the top of the hind paw using a fine needle (30 G) filled with saline (Otsuka Pharmaceutical, Japan) containing 0.3% ethanol (capsaicin or AITC was dissolved in 0.3% ethanol). Mice were gently wrapped in the measurer’s hand and injected in this position. Mice were still very quiet during injection in this position. Their behaviors were recorded using a digital camera (P6000, Nikon, Japan) and analyzed later.

### Statistical Analysis

Data are presented as means ± standard error of mean (S.E.M). Statistical analysis was performed by one way ANOVA followed by a Bonferroni post hoc test or unpaired Student’s *t*-test. p < 0.05 was considered to be significant. Statistical significance is defined as: *, p < 0.05; **, p < 0.01 and ***.

## Acknowledgments

This work was supported by grants to MT from a Grant-in-Aid for Scientific Research from the Ministry of Education, Culture, Sports, Science and Technology in Japan (#15H02501 and #15H05928; Scientific Research on Innovative Areas ‘Thermal Biology’). We thank the Research & Development Laboratory, Dainihon Jochugiku Co., Ltd. (Osaka, Japan) for providing the mosquitoes. We also thank Tomoko Mori and Yumiko Makino (Functional Genomics Facility, National Institute for Basis Biology) for their technical support.

## Author contributions

Author contributions: T. L., S. D., S. A., D. U., S. A. and M. T. designed this study. T. L., S. D., Y. S. and D. U. performed, collected and analyzed data from the experiments. M. T., S. D. and S. A. wrote the manuscript.

